# Anisotropic random walks reveal chemotaxis signaling output in run-reversing bacteria

**DOI:** 10.1101/2020.06.23.163667

**Authors:** Jyot D. Antani, Anita X. Sumali, Tanmay P. Lele, Pushkar P. Lele

## Abstract

The bias for a particular direction of rotation of the flagellar motor is a sensitive readout of chemotaxis signaling, which mediates bacterial migration towards favorable chemical environments. The rotational bias has not been characterized in *Helicobacter pylori*, which limits our understanding of the signaling dynamics. Here, we determined that *H. pylori* swim faster (slower) whenever their flagella rotate counterclockwise (clockwise) by analyzing their hydrodynamic interactions with bounding surfaces. The anisotropy in swimming speeds helped quantify the fraction of the time that the cells swam slower to report the first measurements of the bias. A stochastic model of run-reversals indicated that the anisotropy promotes faster spread compared to isotropic swimmers. The approach further revealed that the diffuse spread of *H. pylori* is likely limited at the physiological temperature due to increased reversal frequencies. Thus, anisotropic run-reversals make it feasible to study signal-response relations in the chemotaxis network in non-model bacterial species.

**Impact Statement:** Anisotropy in run and reversal swimming speeds promotes the spread of *H. pylori* and reveals temperature-dependent behavior of the flagellar switch.

## Introduction

Over half of the human population is colonized by the motile gram-negative bacteria, *H. pylori. H. pylori* infections have been implicated in peptic ulcers as well as non-cardia gastric cancer (1). Infections are promoted by the ability of the bacterium to swim with the aid of helical appendages called flagella (2, 3). The flagella are rotated by transmembrane flagellar motors that repeatedly switch their direction of rotation. Owing to the unipolar location of the left-handed flagellar filaments (4), counterclockwise (CCW) rotation of the motors causes the cell to run with the flagella lagging behind the body – a mode of motility termed as the *pusher* mode. The cell reverses with the body lagging the flagella when the motors rotate clockwise (CW) – a mode of motility termed as the *puller* mode (5). Modulation of the reversals between the two modes enables the cell to undergo chemotaxis (migration towards favorable chemical habitats (6), which promotes pathogenesis. Although the core chemotaxis network proteins and auxiliary proteins are known (7-10), the mechanisms of chemotaxis are not well-understood in *H. pylori* (8).

Switching in the flagellar motor is promoted by the binding of a phosphorylated response regulator, CheY-P, to the flagellar switch. In the canonical chemotaxis network, chemoreceptors sense extracellular ligands and modulate the activity of the chemotaxis kinase. The kinase regulates CheY-P levels to promote switching in the direction of motor rotation (11). Flagellar switching reverses the swimming direction in *H. pylori*. Although prior studies in *H. pylori* have focused on the effects of extracellular ligands on the frequency of cell reversals (12, 13), it has been proposed that the modulation of reversal frequencies by local ligand concentrations alone is not adequate for chemotaxis (14). Instead, the output of the chemotaxis signaling pathway is more accurately quantified by the fraction of the time that the motor rotates CW, termed CW_bias_. The kinase modulates the CW_bias_ to extend runs towards favorable chemical sources (15–17). Dynamic variations in the CW_bias_ sensitively report changes in the kinase activity due to external stimuli as well as due to the internal noise in the network (18, 19).

CW_bias_ has been measured in the model species *Escherichia coli* by tracking the rotation of tethered cells, where a single flagellar filament is attached to a glass surface while the cell freely rotates (15). Alternatively, the bias is determined by sticking the cell to the surface and monitoring the rotation of a probe bead attached to a single filament (20, 21). Such single motor assays have been employed successfully in *E. coli* because the filaments are spaced apart on the cell body. In *H. pylori* however, the filaments are distributed in close proximity to one another at a single pole (22), increasing the likelihood of tethering more than one filament. Tethering of multiple flagella on the same cell eliminates rotational degrees of freedom, inhibiting motor function. Because of the lack of measurements of the CW_bias_, crucial features of the signaling network, such as the dynamic range of signal detection, adaptation mechanisms, and the roles of key chemotaxis-related proteins, remain unknown in *H. pylori* (23, 24).

Here, we report a novel approach to measure the CW_bias_ based on differences in the swimming speeds in the pusher and puller modes of *H. pylori*. We successfully employed this method to quantify the influence of varying temperatures on the interactions between CheY-P molecules and the flagellar motors. With the aid of a quantitative model and simulations, we predict that the random spread of run-reversing bacteria increases with increasing anisotropy in swimming speeds. Our work provides a framework to study chemotaxis signaling and flagellar switch-behavior in *H. pylori*.

## Results

### Swimming speeds are anisotropic

To determine the behavior of flagellar motors in *H. pylori*, we tracked cell motility in the bulk fluid with a phase contrast microscope. The positions of single cells were quantitatively determined from digital videos with the aid of particle tracking (see *Materials and Methods)*. Several cells exhibited reversals in the field of view. A representative cell trajectory at 37°C is shown in **Fig. 1A (**see another example in **Movie S1**). With each reversal, the cell appeared to change from one mode of swimming to the other, although the modes could not be identified (as puller or pusher) because the flagella were not visible. Changes in the swimming modes were distinguished from rotational turns of the cell body – where the swimming mode remains unchanged – by visually inspecting each reversal for each cell. The turn angle between the original direction just before and the new direction just after a reversal (Ø) followed an exponential distribution with a peak ~ 180° (**Fig. 1B**), indicating that cells simply retraced their paths for brief durations following each reversal. The flick of the flagellum that causes turn angles to be distributed ~ 90° in another run-reversing species *Vibrio alginolyticus* (25), is unlikely to occur in *H. pylori*. The distance traveled between any two reversals was identified as a segment and numbered (**Fig. 1A)**. The swimming speeds over 6 consecutive segments are indicated in **Fig. 1C**. The speeds were binned as per the segments, yielding *n+1* bins for *n* reversals. The mean speed from each bin was plotted for all the *n+1* bins (**Fig. 1D**). Mean speeds in alternate bins were anti-correlated: each reversal either decreased or increased the speed. This suggested that the speeds in the two modes were unequal. Such anti-correlation was consistently observed in a large population of cells (n = 250). The distribution of the ratio of their mean speeds in the fast and slow modes is shown in **Fig. 1E**. The speed in the fast mode was ~ 1.5 times the speed in the slow mode.

**Figure 1.**
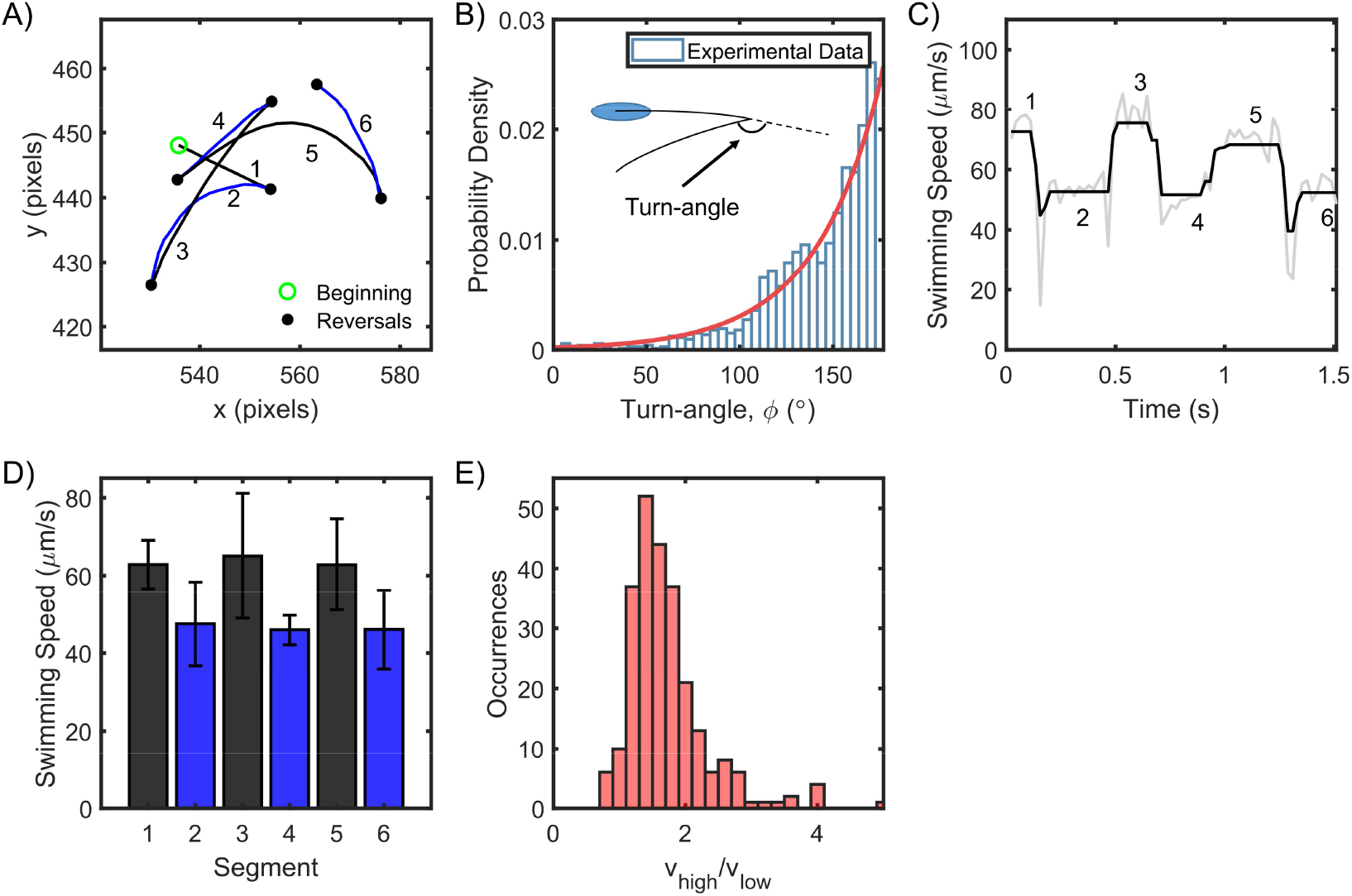
*H. pylori* swim forward and backward at different speeds. **A)** Representative swimming trace of a single bacterium. Each reversal is represented by a filled circle. The beginning of the trajectory is denoted by an open circle. Uninterrupted swimming between two reversals was labeled as a segment and the segments were numbered chronologically. **B)** The turn angles were exponentially-distributed (n = 1653 samples); reversals mostly caused the cells to retrace their movements. **C)** The swimming speed for a single cell over 1.5 s is indicated. The speeds alternated between high and low values with each reversal. Raw data is indicated in gray; filtered data is indicated in black. **D)** The mean speed for each segment is indicated chronologically. Standard deviations are indicated. **E)** The mean speed for the high (low) mode for each cell was calculated by averaging over all its high- (low-) speed segments. The distribution of the ratios of the high and low mean speeds for each cell is indicated. The mean ratio was 1.5 ± 0.4 (n = 250 cells).

### Cell swims faster in the pusher mode

To discriminate between the pusher and puller modes, we exploited the hydrodynamic coupling between swimmers and glass boundaries. Cells that swim very close to an underlying solid boundary exhibit circular trajectories owing to the increased viscous drag on the bottom of the cell and the flagellar filaments. CCW rotation of the left-handed helical filament causes the *pusher* to experience a lateral force that promotes CW circular tracks (**Fig. 2A**, (26, 27)). The situation is reversed when the filaments rotate CW. Thus, it is possible to discriminate between the two modes when a bacterium swims near a surface. We analyzed each cell that swam in circular trajectories near the surface and determined the mean speeds for the two directions. Four sample trajectories are shown in **Fig. 2B**. For each cell, the CW trajectories were always faster relative to the CCW trajectories, indicating that the pusher mode was the faster mode (**Fig. 2C**). This was confirmed over n = 116 cells; the mean ratio of the speeds of the CW trajectories to that of the respective CCW trajectories was ~ 1.6 ± 0.5 (**Fig. 2D**).

**Figure 2.**
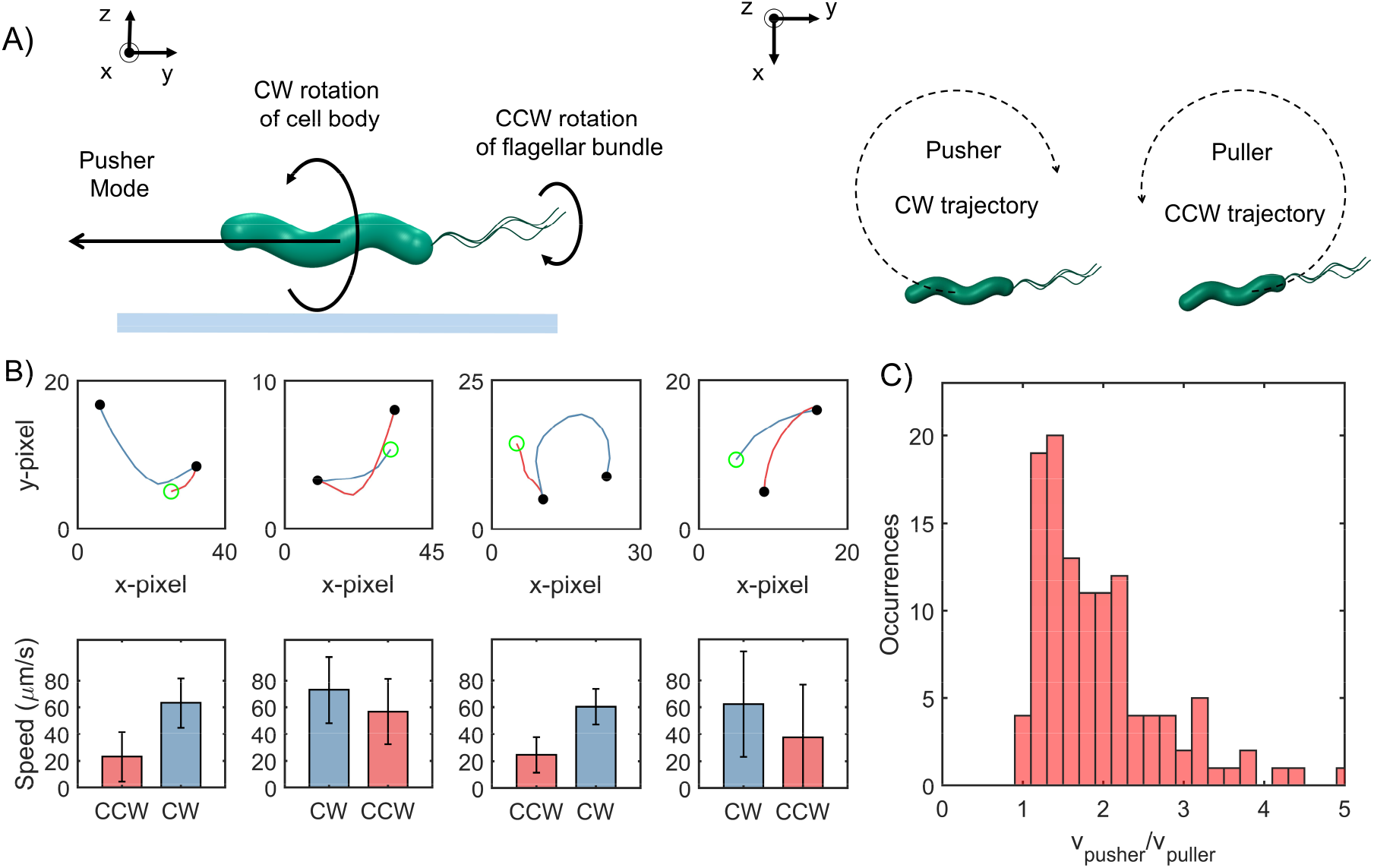
Cells swim faster in the pusher mode. **A)** The viscous drag on the bottom of the cell body and the flagellar filament is higher near an underlying surface (indicated by the blue line in the *left panel*). The drag is lower on the top half of the body and filament. This difference in drag causes a lateral thrust on the cell, giving rise to circular trajectories: CW trajectory in the pusher mode and CCW trajectory in the puller mode (*right panel*). **B)** *Top row:* Blue segments indicate CW trajectories; red segments indicate CCW trajectories. Filled circles indicate reversals; open circle indicates the beginning of the trajectory. *Bottom row:* The corresponding mean speeds and standard errors are indicated for the two trajectories: CW tracks were always faster than CCW tracks. **C)** The distribution of the ratio of the speeds along the pusher and puller modes is indicated (n = 116 cells). The mean ratio = 1.6 ± 0.5.

### Partitioning of swimming speeds enables estimation of CW_bias_

Since *H. pylori* swim in the puller mode when the flagella rotate CW, CW_bias_ could be calculated from the fraction of the time that the cells swam in the slower mode (see *Materials and Methods)*. This method worked for all the cells that reversed at least once in the field of view: the faster and slower modes could be discriminated from each other based on comparisons between the mean speeds before and after a reversal (as shown in **Fig. 1D**). These cells consisted ~81% of the total data. The remaining cells did not reverse under observation; they persisted in a particular direction before exiting the field of view. Hence, these cells were termed as single-mode swimmers. As the mode of swimming could not be readily determined for these cells, those data were grouped into cells that swam near the surface for at least some time and those that did not. In the former group, a majority were identified as pushers based on the direction of their circular trajectories near surfaces, as discussed in **Fig. 2**. About 8% of the cells could not be identified and were excluded from the analysis. The distribution of the bias is shown in **Fig. 3A**. The bias was similar to that observed in *E. coli* (28, 29), suggesting that the basal chemotactic output in the two species is similar. Analysis of the cell tracks obtained at different time-points enabled us to calculate the dynamic variations in the chemotactic output over time (**Fig. 3B**).

**Figure 3.**
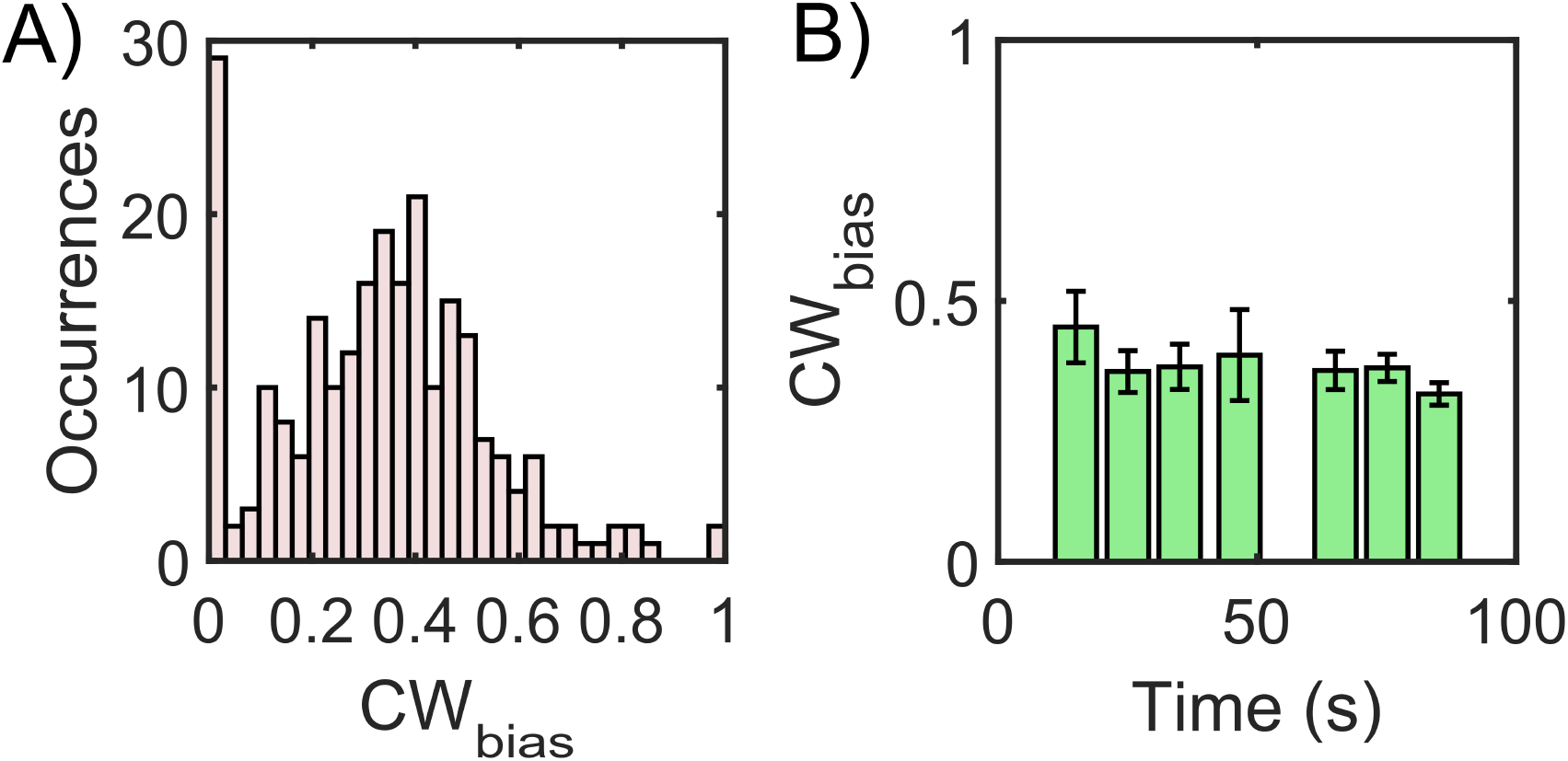
Anisotropic swimming speeds enable quantification of chemotaxis output. **A)** CW_bias_ was determined at 37°C in the absence of chemical stimulants. Cell trajectories with durations of 1 s or more were considered for calculation. The distribution was obtained from n = 240 cells. A Gaussian fit to the switching population (n = 212 cells) yielded CW_bias_ = 0.35 ± 0.23 (mean ± standard deviation). **B)** The variation in CW_bias_ over time was determined by repeating the analysis in **A)** for swimmers recorded over different time points. The time-window for the calculation of each value was ~ 30 s and the sample sizes ranged from n = 11 to 62 cells. Standard error is indicated.

### Effect of thermal stimuli on chemotactic output

The behavior of flagellar motor is strongly influenced by temperature in *E. coli* (30, 32, 33). Considering the similarity in the rotational bias in the two species (**Fig 3A**), we determined whether variations in the temperature had a similar effect in *H. pylori*. We recorded cell motility at different temperatures (see *Materials and Methods)*. The recording began ~ 5 min after each temperature change to provide adequate time for transient processes to stabilize. The mean pusher and puller speeds trended upwards with the temperature (**Fig. 4A**, left panel), presumably through modulation of proton translocation kinetics that power the motor (33). The ratio of the speeds in the two modes appeared to be independent of the temperature (**Fig. 4A**, right panel).

**Figure 4.**
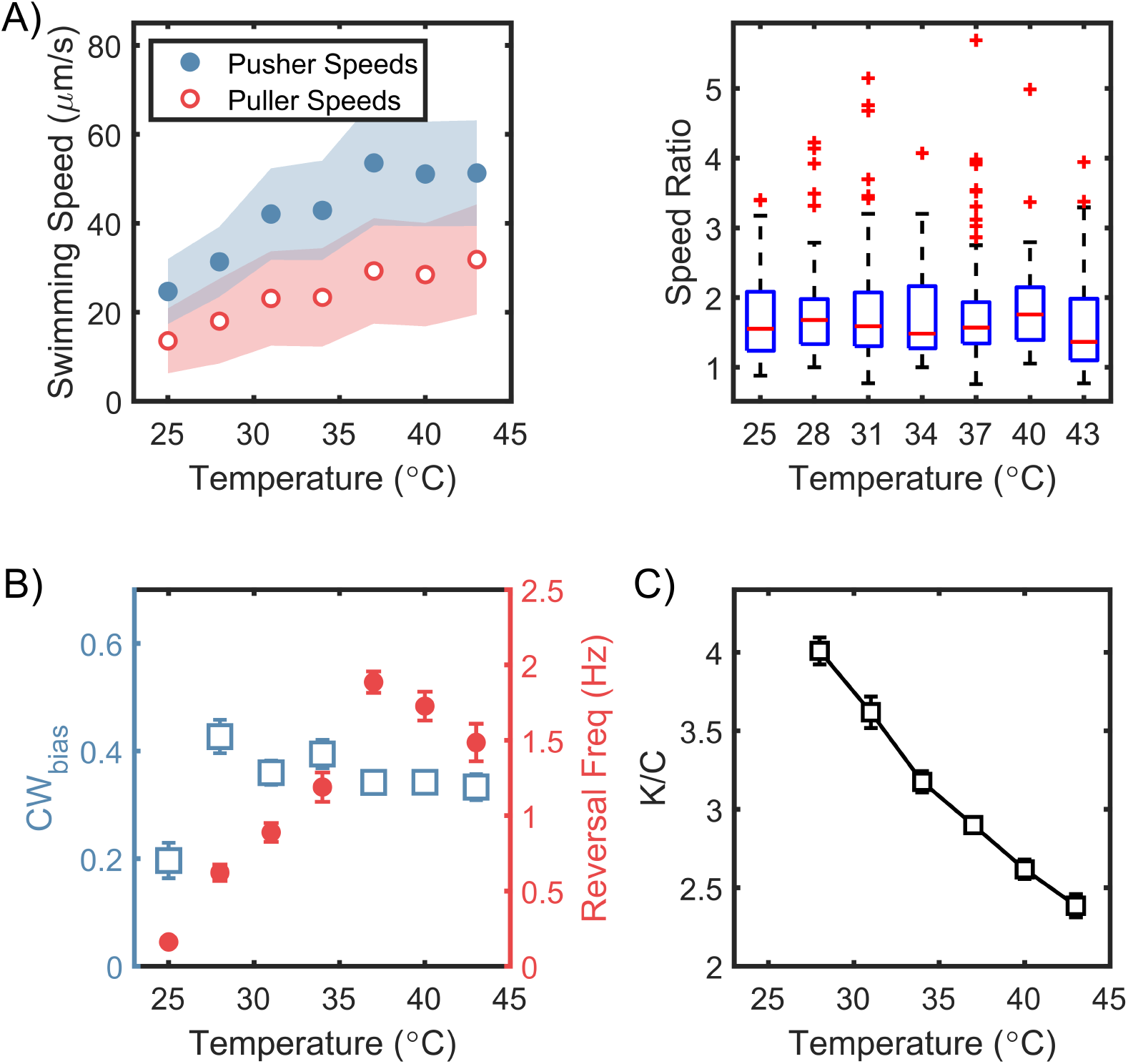
Steady-state chemotactic output is independent of temperature. **A) *Left:*** Swimming speeds for each mode are plotted (mean ± standard deviation) for different temperatures. The speeds increased with temperature till 37°C, after which they plateaued. The shaded regions indicate standard deviation. ***Right:*** The ratios of the pusher and puller speeds are indicated. The ratios appear independent of the temperature. **B)** Mean CW_bias_ (open squares) and mean reversal frequencies (filled circles) are plotted over a range of temperatures. The switching frequency was at a maximum at the physiological temperature (37°C), and decreased at higher and lower temperatures. The CW_bias_ increased with the temperature and plateaued above 30°C. The mean values are indicated with standard error. Each data-point was averaged over n ≥ 50 cells. **C)** The estimated ratio of the CheY-P dissociation constant (*K*) and the intracellular CheY-P concentrations (*C*) is indicated as a function of the temperature. The ratios were calculated from the data in **B)** following a previous approach (30). The number of binding sites for CheY-P in *H. pylori* ~ 43 was estimated from the relative sizes of the flagellar C-ring (see *Supplementary Information* and ref. (22)). The ratio of the dissociation constants for the CCW and the CW motor conformations was assumed to be similar to that in *E. coli* (~ 4.7 from (31)).

The frequency of reversals increased steadily with temperature up to 37°C. Interestingly, the steady-state CW_bias_ varied weakly with the temperature (**Fig. 4B**); thus, the effect of temperature on the steady-state chemotaxis output is relatively minor. Flagellar switching can be described well by a two-state model, where the binding of phosphorylated CheY (CheY-P) to the flagellar switch stabilizes the CW conformation (30). In the absence of CheY-P, the probability of observing CW rotation in an otherwise CCW-rotating motor decreases with increasing temperatures (32). Thus, the relative insensitivity of the rotational bias in **Fig. 4B** suggests that the dissociation constant for CheY-P/switch interactions likely decreases with rising temperatures. Following the thermodynamic analysis of Turner and co-workers (30), we calculated the normalized estimates of the dissociation constant from our data, as shown in **Fig. 4C** (see *Supplementary Information* for details). It is reasonable to extend the framework to *H. pylori* considering the similarities between its chemotaxis output with that of *E. coli* (**Fig 3A**). For similar CheY-P levels as *E. coli*, we estimate the dissociation constant to be ~ 9 μM at 37°C.

### Speed anisotropy promotes diffusion

Bacterial motion is uncorrelated over long times and large length-scales in the absence of a signal. The random walk with exponentially distributed run-times has been quantified with an effective diffusion coefficient for a run-tumble model such as *E. coli* (14, 34, 35). To model the diffusion of *H. pylori*, we adopted a stochastic reversal model for isotropic run-reversals (36) and modified it to incorporate anisotropic swimming speeds (see *Supplementary Information* for details). Briefly, the velocities of a bacterium that swims at *v*_0_ μm/s in its slower mode was expressed as: ***v***(*t*) = *v*_0_ *h*(*t*) [1 + *aH*(*h*)]*e*(*t*). The direction of swimming was described by the function *h*(*t*), which alternated between +1 and −1 with each reversal (**Fig. 5A**). A Heaviside function, *H*(*h*) and the anisotropy parameter, *a*, characterized the magnitudes of the speeds in the two directions: *v*_0_ and *v*_0_(1 + *a*). The CW_bias_ was assumed to be constant (= 0.5) to simplify the model.

**Figure 5.**
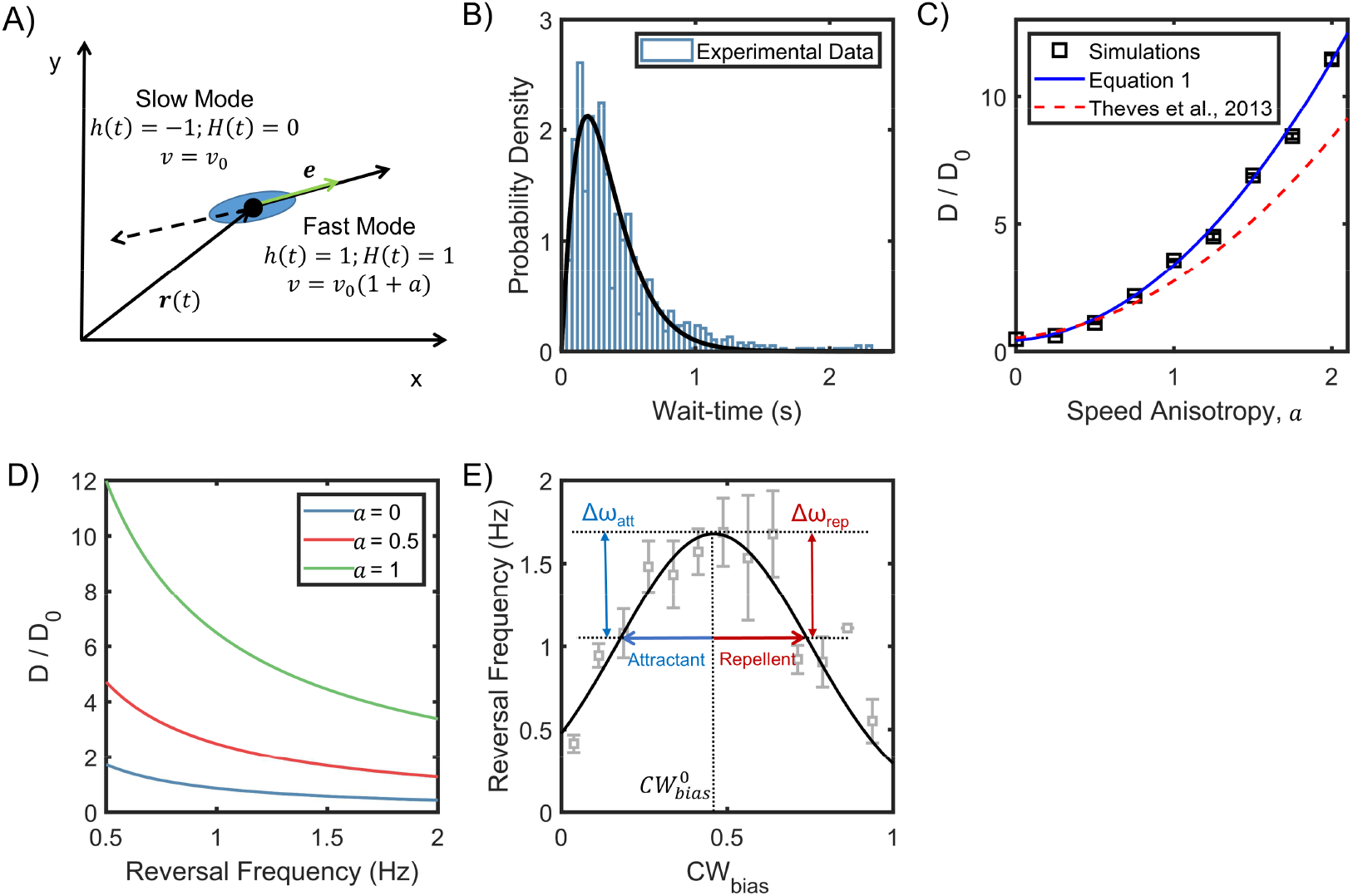
Anisotropic random walks in a run-reversing bacterium. **A)** Cell swims at *v*_0_ μm/s in the puller (slower) mode, and at *v*_0_(1 + *a*) μm/s in the pusher (faster) mode. The isotropic case is described by *a* = 0, where the run and reverse speeds are equal. Cell alignment is described by the unit vector ***e***. **B)** Experimentally observed wait-time intervals for runs and reversals obey a Gamma distribution (n = 983 samples): the shape and scale parameters were *k* = 1.94 ± 0.15 and *θ* = 0.21 ± 0.02, respectively. **C)** The diffusion coefficients predicted from equation 1 are indicated as a function of the anisotropy in speeds (blue curve). Predictions from an alternative model for anisotropic swimming is shown by the dotted curve (37). Symbols indicate coefficients calculated from simulation runs (see *Supplementary Information*). The parameters were based on experimental measurements: mean wait-time = 0.5 s, α = 0.86, and *v*_0_ = 25 μm/s. *D_θ_* = 0.02 s^−1^ from (36). Diffusion coefficients have been non-dimensionalized with 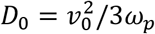 (34), where *ω_p_* is the reversal frequency at the physiological temperature (**Fig. 4B**). **D)** Predicted diffusivity is indicated over a range of typical reversal frequencies. Here, α = 0.86 and *D_θ_* = 0.02 s^−1^. **E)** The symbols indicate the experimental relationship between the reversal frequency and CW_bias_ in *E. coli* (38), and the black curve indicates a fit. The blue and red arrows indicate the effect of attractants and repellents on the CW_bias_, respectively. The corresponding changes in the reversal frequency are similar (Δω_att_ ~ Δω_rep_).

The deviation of the cell from a straight line during a run (or reversal) occurred due to rotational diffusion, described by 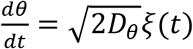. White noise characteristics were 〈*ξ*(*t*)〉 = 0, and 〈*ξ*(*t*)*ξ*(*t+τ*)〉 = *δ*(*τ*), where *D_θ_* is the rotational diffusion coefficient. Another randomizer of the bacterial walk is the turn angle, Ø, which is the angle between the original direction just before and the new direction just after a reversal. The turn angle is likely influenced by kinematic properties: cell shape, filament bundling dynamics, and the flexibility of the flagellar hook. Next, the wait-times distributions were experimentally characterized (**Fig. 5B**), and the fitted shape parameters (*k, θ*) were subsequently employed to describe the reversal dynamics as discussed in the *Supplementary Information*. This enabled us to derive an expression for the asymptotic diffusion coefficient from the velocity correlation over long-times:

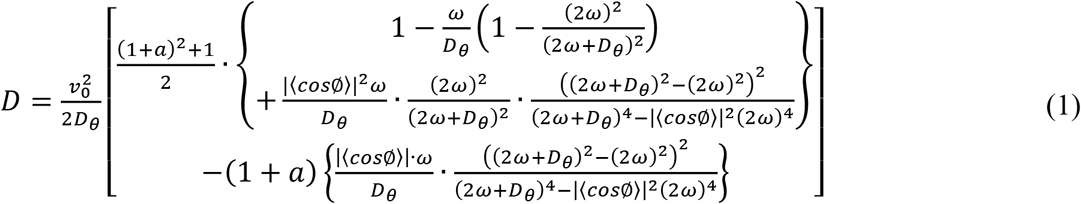

The reversal frequency is indicated by ω. The expression correctly reduces to that for the isotropic swimmer (36), for α (= |〈*cos*Ø〉|) = 1, and *a* = 0.

The diffusion coefficients increased with the anisotropy, *a*, as shown in **Fig. 5C**. Model predictions were validated via comparisons with the diffusion coefficients calculated from stochastic simulations of 1000 cells that underwent run-reversals (see *Supplementary Information*). Thus, anisotropic run-reversers (*a* ≠ 0) are likely to spread faster than isotropic run-reversers (*a* = 0). The diffusion coefficient was inversely dependent on the run-reversal frequency (**Fig. 5D**). Since the reversal frequencies reach a maximum at 37°C (**Fig. 4B**), it is possible that cells at physiological temperatures spread slower in a niche over long-times, providing more time for cells to adhere to surfaces.

## Discussion

The response of the chemotaxis signaling pathway to external stimuli is conventionally measured by determining the rotational bias (15, 17, 39, 40). This is done by sticking probes to individual flagellar motors and tracking their rotation. Alternately, the signaling output has been determined via Förster resonance energy transfer-based measurements of *in vivo* enzymatic reactions (41). Neither of these approaches has been realized in *H. pylori:* the close proximity of flagella (22) increases the chances of multiple motors being tethered simultaneously and stalling, and functional fusions of chemotaxis proteins with FRET pairs are lacking. Our approach circumvents these limitations and relies on facile light microscopy to determine the rotational bias as a measure of the signaling output at a single cell and population level. The distribution of the rotational bias was similar to that observed in *E. coli* (28, 29), suggesting that the basal chemotaxis activity in the two species is similar.

Recent studies on chemotaxis in *H. pylori* focused on the effect of chemical stimulants on the reversal frequency (12, 13, 42). Because diffusion scales inversely with the reversal frequency, increases in frequency might help a cell linger in a niche. However, mere dependence of the reversal frequencies on the local temperatures alone is not adequate for thermotaxis; such dependencies would yield uniform distributions of cells even in a varying thermal field. Importantly, the reversal frequencies can be similar at low and high kinase activities (38), as shown in **Fig. 5E**. Thus, an attractant and a repellent might induce similar changes in the reversal frequency despite opposing effects on the kinase activity. Hence, categorization of chemoeffectors as attractants or repellents is more reliably achieved by studying their effects on the CW_bias_.

To prevent the cell from tumbling during a reversal, all the flagellar motors in a single cell of *H. pylori* must switch synchronously from one direction to the other. Indeed, no tumbles were observed in our experiments. The most frequent turn angles were ~ 180°, which confirmed that the cells retraced their paths following a reversal – this would not have been the case if only a fraction of the motors switched to the opposite direction. This makes our approach feasible for determining the CW_bias_ for an individual cell from its swimming speeds – which reflects the collective action of all the motors – rather than sampling individual motors. Interestingly, switching in the direction of motor rotation is stochastic (in *E. coli*), and is well approximated by a two-state Poisson process with exponential wait-time intervals (43). This is likely true for *H. pylori* as well. The actual wait-time intervals for the run-reversals in *H. pylori* obeyed a Gammadistribution (**Fig. 5B**), but this mostly reflects contributions of the torsional relaxation dynamics of the flexible flagellar hook that connects the flagellar motor and the flagellar filament (44). How are such multiple stochastic switchers coupled in *H. pylori*? One possibility is that the flagellar switch in *H. pylori* is ultrasensitive to small fluctuations in CheY-P levels, similar to the switch in *E. coli* (18). The close proximity of the multiple motors at a single pole in *H. pylori* also means that the local concentration of CheY-P in the vicinity of each flagellar switch is similar. This increases the probability of concerted switching in all the motors.

Our simulations and model indicate that run-reversing bacteria that undergo anisotropic random walks diffuse faster than isotropic run-reversers. Anisotropic run-reversals in some species have been observed near bounding surfaces (37, 45). In *H. pylori*, we observed anisotropic speeds in some cells even at a separation of ~ 200 μm from any bounding surfaces (**Fig. S4**). Therefore, the anisotropy is unlikely to be a surface-effect. The effect could be due to differences in the flagellar shapes and bundling or the swimming gait in the pusher and puller modes. However, it is more likely that the anisotropy arises due to the differences in the CW and CCW flagellar rotational speeds, as is the case with *E. coli* – which run and tumble – and *Caulobacter cresecentus* (21, 46) – which exhibit isotropic run-reversals (**Table S2**). The rotational direction-dependent differences in motors speeds in the two species depend on the external viscous loads (21, 46). It is possible therefore, that the anisotropy in *H. pylori* is also load-dependent; vanishing for longer filament lengths in highly-viscous microenvironments or for very short filaments. The anisotropy is further expected to depend sensitively on the expression of the flagellar genes, which is modulated by environmental conditions (47). The anisotropy was prominently observable in our work with a careful control of experimental conditions (*Materials and Methods*).

We anticipate that the analysis described in this work will help identify the molecular determinants of chemotaxis adaptation in *H. pylori*, and explain how the receptors retain sensitivity to ligands over a wide range of concentrations. The method is extensible to any run-reversing species that exhibit anisotropic swimming speeds, paving the way to study signaling dynamics in other run-reversing bacterial species.

## Materials and Methods

### Strains and cell culturing

All the work was done with *H. pylori* PMSS1. Cultures of microaerophilic *H. pylori* were grown in an incubator with controlled temperature and CO_2_–environment (Benchmark Incu-Shaker Mini CO_2_). The incubator was maintained at 10% CO_2_, 37°C. Fresh colonies were streaked out before each experiment on Columbia horse blood agar (Thermo Scientific™) plates supplemented with Polymyxin-B (Alfa Aesar), Vancomycin (Sigma Aldrich), and 5% w/v defibrinated horse blood (Hemostat Laboratories).

Multiple colonies from the agar plates were picked with the aid of sterilized cotton-tipped applicators and inoculated in 5 mL of BB10 (90% Brucella Broth, Millipore Sigma + 10% Fetal Bovine Serum, Gibco™) to grow overnight cultures. No antibiotics were added to the liquid cultures as per established protocols (4, 13, 42). The overnight cultures were grown for ~16 hours till an OD_600_~0.25-0.5. The overnight culture was then diluted to OD_600_~0.1 by adding fresh BB10, and grown in the shaker incubator set at 170 rpm under 10% CO_2_ and at 37°C.

The day cultures were grown to an OD_600_~0.125-0.15 and subsequently diluted in a motility buffer (MB, 0.01 M phosphate buffer, 0.067 M NaCl, and 0.1mM EDTA, pH~7.0) prior to imaging. The dilution was ~6-7% v/v (BB10/MB). Cells remained motile in MB for over an hour.

### Motility assays

A culture-dish provided as part of a micro-environmental temperature control system (Delta T™ system, Bioptechs Inc.) was used for measuring motility. Cells were imaged on a Nikon Optiphot with a 10X objective, which was installed inside a temperature-controlled chamber (ETS Model 5472, Electro-Tech Systems, Inc). Videos were recorded with a CCD camera (IDS model UI-3240LE) at 45 fps.

### Temperature control

The temperature control chamber (ETS Model 5472, Electro-Tech Systems, Inc) enabled precise modulation of the temperature during cell visualization. After each change in the temperature, ~ 5 – 10 min were provided before recording videos.

### Visualization of flagella

A high numerical phase objective (100x, NA 1.49 mounted on a Nikon Eclipse Ti-E microscope) was used for visualizing flagellar bundles in swimmers. Videos were recorded at ~ 100 fps.

### Data analysis

The dilution in MB yielded a cell density that was appropriate for quantifying cell movements with the aid of particle-tracking methods (20). All the videos were analyzed with custom-written MATLAB codes based on centroid-detection-based particle-tracking routines (48). The experimental data presented in the figures were from 5 or more biological and multiple technical replicates, except Fig 4. The data in Fig 4 were obtained from a single biological and multiple technical replicates.

### CW_bias_ calculations

For each cell, the swimming speeds alternated between high and low values with every reversal. A visual scan of the recorded movie was made with ImageJ (NIH) to confirm the number of reversals. Since the speeds in the two modes were distinct, a bimodal distribution was obtained per cell. This enabled us to select a unique threshold for each cell to bin the speeds. For each cell n + 1 bins were obtained for n reversals. A reversal changes the mode of motility between the pusher and the puller mode. On the other hand, a 180° turn by the cell maintains the same mode. Each reversal was therefore confirmed visually to distinguish between reversals and the rare 180° turns by the cells. This process was repeated for hundreds of cells.

To determine the CW_bias_, cells that were observed for at least 0.5 s were retained for analysis. CW_bias_ was calculated as the fraction of the time that a cell swam in the puller (slower) mode, which corresponds to CW rotation of the filament. To do this, the number of frames in which the i^th^ cell swam in the puller mode, 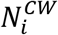, was divided by the total frames over which the cell was observed, *N_i_*, to yield: 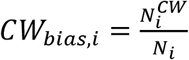

The error associated with the calculation of *CW_bias,i_* values decreases with increasing *N_i_*. But, different cells were observed for different durations; hence the *CW_bias,i_* values were allocated weights that corresponded to their respective durations: 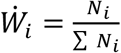. Mean bias was determined as:

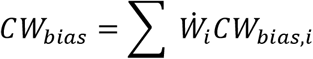

Reversal frequency was determined in a similar manner.

## Acknowledgments

We thank Professor Karen Ottemann for the PMSS1 strains, and Professor Michael Manson and Rachit Gupta for comments. PPL acknowledges funding from the Cancer Prevention and Research Institute of Texas (RP170805) and the National Institute of General Medical Sciences United States (R01-GM123085).

## Author contributions

JDA and PPL designed the work and wrote the paper; JDA carried out the experiments; JDA and AXS analyzed the data; TPL and PPL developed the model; PPL developed the simulations and experimental setups.

